# Integrative single cell analysis of CD8+ T-cells across early and advanced oral cancers reveals signatures of anti-tumour activity

**DOI:** 10.64898/2026.09.02.748952

**Authors:** Constance H Li, Hong Sheng Quah, Camille Arcinas, Lakshmi Alagappan, Lisda Suteja, Hariraman Bhuvaneswari, Hui Sun Leong, Fui Teen Chong, Darren Toh, Sophia Wong, Subhra Kumar Biswas, Cliburn Chan, N Gopalakrishna Iyer

## Abstract

Tumour-targeting CD8^⁺^ T cells drive responses to every major form of cancer immunotherapy. Identifying them, however, remains an unsolved problem in solid tumours. The antigens they recognize are rarely defined and almost never shared between patients. We profiled 51,459 CD8+ T cells by paired single-cell RNA and T-cell receptor sequencing across 28 samples from 17 HPV-negative oral cancers spanning primary tumours, draining lymph nodes, metastases, and pembrolizumab-treated recurrences. We found that clonotypes that were expanded and shared across anatomical sites and timepoints were enriched within tumours and progressively selected over disease evolution and checkpoint blockade. Designating these shared-expanded clones as putative tumour-targeting cells, we trained a machine learning classifier that identifies them from transcriptome data alone. This 108-feature random forest signature recapitulated programmes of tumour reactivity and generalized to an integrated atlas of 89,318 CD8+ T cells from independent cohorts, showing progressive enrichment from normal to malignant tissue, and localized to tumour-proximal niches in spatial transcriptomics. By demonstrating that clonal behaviour across space and time encodes tumour reactivity in the transcriptome, this work establishes a generalizable framework for mapping tumour-engaged immunity without knowledge of the underlying antigen.

## Introduction

Tumour-targeting cytotoxic CD8^⁺^ T cells are the critical effectors that underpin the success of modern cancer immunotherapies. Their recognition, recruitment, and reinvigoration form the basis of the most clinically impactful cancer immunotherapy modalities, including immune checkpoint blockade (ICB), bispecific antibodies, engineered T cell therapies, and adoptive transfer of tumour-infiltrating lymphocytes (TILs).^1–4^. Despite their central role, reliably distinguishing the T cells that recognize and kill tumour cells from the broader infiltrating population remains an unsolved problem in solid tumours^5,6^. The gold standard for defining tumour specificity is direct demonstration that a T cell receptor (TCR) binds its cognate tumour-derived antigen^7,8^.This approach is highly effective in rare contexts where tumour antigens are clearly defined, such as viral oncoproteins in HPV-positive cancers, tumour-associated antigens like the melanoma-associated antigens MART-1 or gp100, or neoantigens created by recurrent hotspot mutations (*eg*. KRAS G12D, PIK3CA H1047R)^9–11^. However, these contexts represent the exception rather than the rule: immune editing systematically eliminates specific T-cells to progressively sculpt the tumour-immune microenvironment towards evasion, and renders the identity of tumour-reactive clones largely invisible to antigen-dependent identification strategies^12,13^.

The problem is compounded in tumour types where the antigen landscape itself is poorly defined, heterogeneous, or suppressed. Most solid tumours, including the majority of HPV-negative head and neck squamous cell carcinomas (HNSCC) lack universal, shared, or easily detectable immunogenic antigens^14,15^. Tumour-derived neoantigens are infrequent, highly personalized and rarely conserved across patients, making antigen-resolved methods impractical at scale. Computational approaches to neoantigen prediction have offered a partial remedy, but carry substantial false positive rates and reduced reliability across less common HLA haplotypes^16,17^. Deprived of dependable antigen-based strategies, the field has instead relied on indirect features such as markers of T cell exhaustion, tissue residency, clonal expansion, or cytotoxic differentiation to infer tumour specificity, though each alone provides an incomplete picture^18^. A reliable and generalizable method to identify tumour-targeting CD8^⁺^ T cells in solid tumours therefore remains a major unmet need.

Recent advances in single-cell RNA sequencing (scRNA-seq) and paired T cell receptor sequencing (scTCR-seq) offer a transformative solution. These approaches enable high-resolution, unbiased profiling of intratumoural T cells, capturing their transcriptional states, activation and dysfunction programs, and clonal architecture. Integrating scRNA-seq with scTCR-seq allows reconstruction of T cell differentiation trajectories and tracking of shared clonotypes across tissue compartments, revealing how tumour-reactive clones emerge, expand, become dysfunctional, or respond to therapy^19–22^. Importantly, these technologies permit multidimensional inference of tumour specificity combining phenotype, clonal expansion, and activation status.

The surgical management of HPV-negative HNSCC provides a unique opportunity for comprehensive tissue sampling across both space and time. Here, we profile tumour samples reflecting different tissue contexts across varying disease stages in HPV-negative oral cavity squamous cell cancer (OSCC) patients, spanning primary tumours, involved and uninvolved tumour-draining lymph nodes, and tissues obtained at recurrence or following ICI salvage procedures. Integrating scRNA-seq and scTCR-seq, we delineate the molecular and clonal hallmarks of tumour-targeting CD8^⁺^ T cells, reconstruct the progression of tumour-reactive clones across compartments and treatment exposures, and derive a robust, antigen-independent framework for identifying tumour-targeting CD8^⁺^ TILs in solid tumours.

## Results

### Single-cell profiling of CD8^⁺^ T cells across the oral cavity cancer disease spectrum

We collected 28 tissue samples from 17 oral cavity cancer patients reflecting early, intermediate and late disease (**Supplementary Table 1**), and performed scRNAseq and scTCRseq analyses of CD8^+^ T cells across this spectrum of disease progression (**Figure 1A & 1B**). Nine patients had early disease, defined as a primary oral cavity tumour with no clinical or pathological evidence of lymph node involvement; we obtained primary tumour tissue samples (Pri) from all nine patients and matched, pathologically normal tumour-draining lymph node tissue (TDLN) from three patients (HN307, HN309, and HN325). The intermediate disease group comprised four patients who presented with nodal metastatic disease. We obtained primary tumour tissues for all four intermediate disease patients and a paired nodal metastasis (LNMet) sample from patient HN290. The late disease group included four patients who developed distant metastases, allowing analysis of both primary and recurrent metastatic tumours (Rec). Furthermore, HN338 and HN386 received pembrolizumab as single agent treatment for the respective recurrences. For these tumours, additional on-treatment samples were collected one week after treatment initiation (Rec-pICB), allowing direct assessment of early CD8^+^ T cell responses to PD1 blockade.

**Figure 1.**
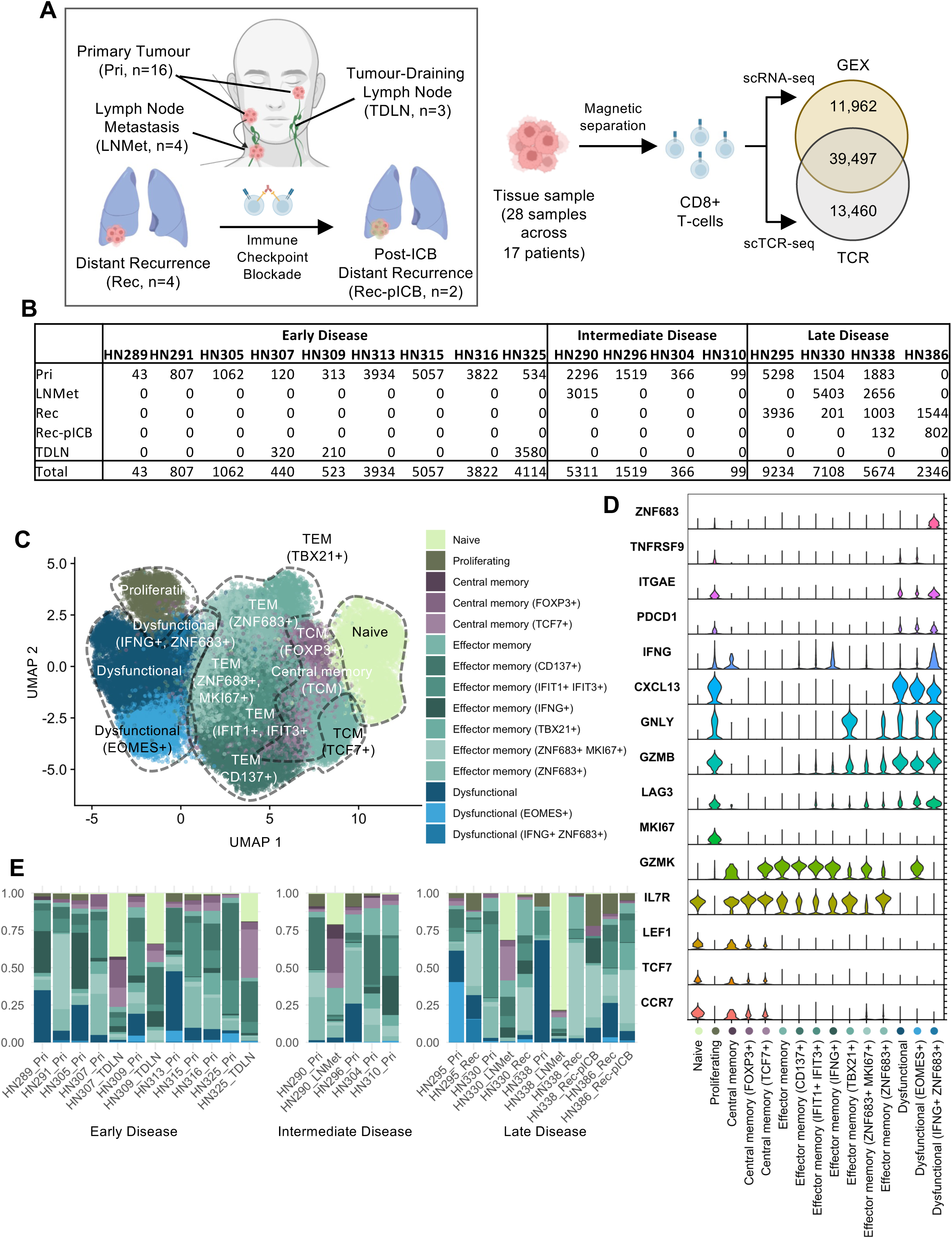
Single-cell sequencing reveals CD8+ T-cell populations across stages of HNSCC. A. Study data include multiple sites spanning oral cavity cancer disease stages and treatment. Samples were magnetically enriched for CD8+ cell populations, which were profiled by scRNA-seq and scTCR-seq. B. scRNA-seq cell counts after quality filtering per sample and summarized across patients. C. Fine-resolution CD8+ cell clusters following a 2-stage clustering approach. Initial broad cluster regions are indicated by dotted lines. D. Marker genes used to annotate fine-resolution clusters. Both known marker genes and differentially expressed genes were used. E. Cell cluster proportional breakdown with colour indicating cluster annotations consistent with legend in (C)

We enriched CD8^+^ T cells by magnetic-activated cell sorting (MACS) and profiled these by scRNA-seq and scTCR-seq. After filtering gene expression data for quality, we obtained a high-confidence population of 51,459 CD8^+^ T cells across the 28 samples. After data processing, normalization and integration, the harmonized dataset showed no residual structure attributable to covariables including patient ID and tissue type (**Supplementary Figure 1A**). Clustering revealed 5 major cell clusters representing canonical CD8^+^ T cell states of naïve, central memory, effector memory, proliferating and dysfunctional (**Supplementary Figure 1B**). For each of these populations, we further subclustered to refine the resolution of heterogenous CD8^+^ T cell states. This approach identified seven effector memory cell sub-clusters, three central memory cell sub-clusters, and three dysfunctional cell sub-clusters, but did not break down naïve or proliferating compartments into further sub-populations (**Figure 1C**).

Effector memory sub-clusters included populations characterized by elevated markers of active TCR signalling (*CD137^+^/IFNG^+^*) indicating acute antigen engagement and cytotoxic potential^23^, tissue-residency (*ZNF683^+^/TBX21^+^*) supporting local immune surveillance and long-term tumour control, and interferon-induced response genes (*IFIT1^+^/IFIT3^+^*) suggesting enhanced anti-viral and anti-tumour immune responses (**Figure 1D**, **Supplementary Table 2**). Central memory clusters included *TCF7^+^* stem-like cells with self-renewal potential^24^. Dysfunctional CD8^+^ cells comprised clusters with intermediary (*IFNG^+^ZNF683^+^*) and late dysfunctional characteristics (*EOMES^+^*), highlighting populations that may have the potential for therapeutic reinvigoration for effective tumour elimination (**Figure 1D**, **Supplementary Table 2**).

Stratification of cell populations by disease category and tissue type revealed specific patterns of enrichment. Nodal tissues (including TDLN and LNMet) contained higher proportions of naïve and central memory CD8^+^ T cells, including TCF7^+^ stem-like populations, consistent with the role of lymph nodes in T cell priming and maintenance of stem-like memory reservoir regardless of tumours involvement (**Figure 1E**, **Supplementary Figure 1D**). In contrast, primary tumours across disease categories had increased frequencies of effector memory, dysfunctional and proliferating CD8^+^ T cells, consistent with local antigen-driven activation and exhaustion in the tumour microenvironment. Altogether, these diverse populations define the continuum of CD8^+^ T cell states within the tumour and draining lymph nodes.

### scTCRseq reveals T-cell expansion dynamics through differentiation

Paired scRNA-seq and scTCR-seq profiling links transcriptional state to clonal identity, enabling systematic reconstruction of T cell clonal dynamics across disease progression. Our data includes paired scRNA-seq and scTCR-seq profiles for 39, 497 cells (**Figure 1A**). The scTCR-seq data captured varying numbers of cells across samples (mean 1,857, range 61-5,346) and a similarly wide diversity of unique TCR clones (mean 957, range 55-3,597; **Supplementary Table 3**) per sample. On average, unique clones made up 61.1% (range 24-98.7%) of each sample’s CD8+ TCR repertoire (**Figure 2A**). Lymph node tissues displayed significantly higher TCR clonal diversity when compared with tumour tissues, regardless of nodal involvement by metastatic disease (p=0.017, effect = 23.5% (4.2-49.4% 95%CI), Wilcoxon test; **Figure 2B**). Notably, no significant difference was observed in clonal diversity between LNMet and TDLN.

**Figure 2.**
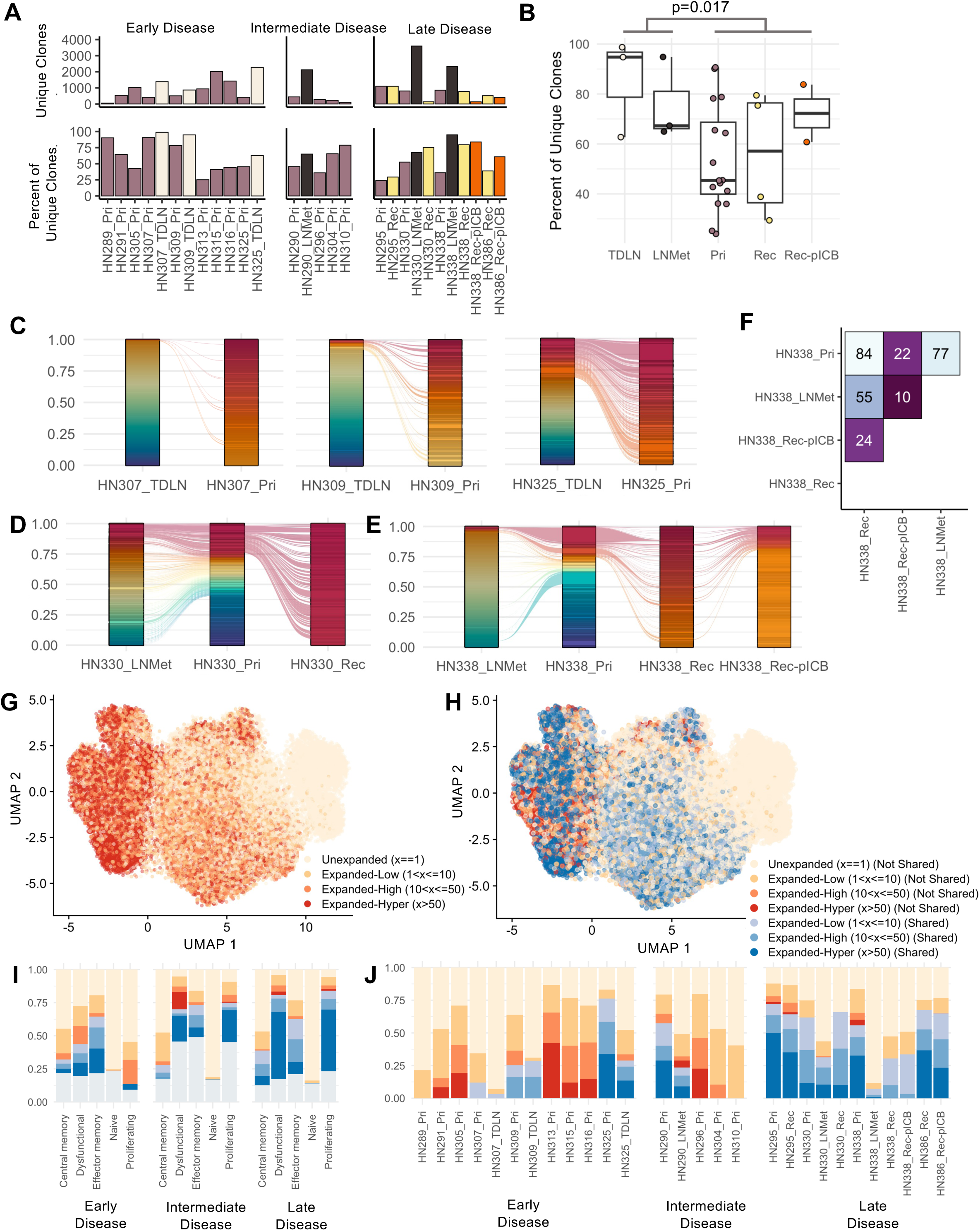
CD8+ T cell clone tracking shows expansion dynamics through tissue type and disease stage. A. Per-sample clonotype richness shown as the number of unique clonotypes detected (top) and the proportion of TCR-recovered cells belonging to unique clonotypes (bottom). Colours denote tissue type, with brown=Pri; white=TDLN; black=LN-Met; yellow=Rec; orange=Rec-pICB. B. Clonotype richness, expressed as the proportion of unique clonotypes per sample, is significantly higher in lymph node tissues than tumour tissues. Box plots show median and IQR; p-value reflects Wilcoxon rank-sum test comparing lymph node (TDLN and LNMet) versus tumour tissues (Pri, Rec, and Rec-pICB). C. Alluvial plots of TCR clonal overlap between primary tumour and matched tumour-draining lymph node (TDLN) in three early-stage patients (HN307, left; HN309, centre; HN325, right). Each ribbon represents a unique clonotype; ribbon width reflects its proportional contribution to the sample TCR repertoire. Ribbons spanning both columns indicate clonotypes shared between compartments. D-E. TCR clonal overlap in late disease patient samples, between (D) lymph node metastasis, primary tumour, and recurrent tumour in HN330 and (E) lymph node metastasis, primary tumour, recurrent tumour and post-pembrolizumab recurrent tumour in HN3308. F. Absolute counts of shared clonotypes between pairs of samples for HN338 G. Clonotype expansion state overlaid on scRNA-seq UMAP. Cells are coloured by clonal expansion category derived from scTCR-seq: unexpanded (x=1), lowly expanded (1<x≤10), highly expanded (10<x≤50), and hyperexpanded (x>50), where x denotes the number of cells sharing an identical paired CDR3 nucleotide sequence. H. Clonotype expansion and sharing as defined by scTCR-seq overlaid on scRNA-seq-derived UMAP. I-J. Proportions of clonotype expansion-sharing by (I) coarse cell cluster and (J) sample across disease stage. Grey NA values indicate cells profiled by scRNA-seq but not scTCR-seq.

We next examined clonal overlap across samples, asking whether expanded clonotypes were shared within or between individuals. As expected, clonotype sharing was largely restricted within individuals, with only rare instances of overlapping clones detected in more than one individual (**Supplementary Table 3, Supplementary Figure 2A**). We subsequently focused on intra-patient clone sharing between samples derived from the same individual. For three early disease patients, we compared shared clones between primary tumour and matched TDLN, finding considerable variability in the number of shared clones (11 to 154 shared clones; **Figure 2C**, **Supplementary Figure 2B**). We extended this comparison to all primary tumour-lymph node pairs (including intermediate and late diseases) and found similar variability across local metastatic disease (**Supplementary Table 3**). However, across all disease stages, shared clones consistently represented a greater proportion of the clonal repertoire in primary tumours than matched lymph node tissues (Paired Mann-Whitney test p = 0.031, **Supplementary Figure 2C**).

Longitudinal tumour sampling in late disease patients presented a unique opportunity to trace T-cell clonal dynamics over time and clinical intervention. Patient HN330 had a primary tumour and synchronous LNMet, then experienced a distant recurrence nine months after surgery with lung and dermal metastases; the surgically accessible dermal metastasis was profiled by single sell sequencing. HN330’s primary tumour shared a more diverse set of clones with the LNMet than the subsequent recurrent tumour (**Figure 2D, Supplementary Figure 2D-E**): the primary tumour and LNMet shared 258 clones, accounting for 56% of primary tumour T cells and 30% of LNMet T-cell repertoire. In contrast, only 62 clones were shared between the primary (22%) and recurrent (60%) tumours. A further 45 clones were shared exclusively between the LNMet (0.3%) and recurrent (4%) tumours. The three sites shared 38 persistent overlapping clones dominating the repertoire of primary tumour (19%), LNMet (10%) and recurrence (39%; **Supplementary Table 3**, **Supplementary Figure 2E**).

Patient HN338 also had primary tumour and synchronous LNMet, with metastases to the lung, bones and chest wall six months after surgery. Biopsy samples of the chest wall metastatic tumour were taken before (Rec sample) and two weeks after Pembrolizumab initiation (Rec-pICB sample). We observed similar dynamics of clonal sharing as in patient HN330, with 77 overlapping clones between the primary (40%) and LNMet (5%) tumours *versus* 84 overlapping clones between the primary (25%) and pre-ICB recurrent (15%) tumours (**Figure 2E-F; Supplementary Figure 2F**). Interestingly, introduction of ICB altered this landscape, where the on-treatment recurrent tumour only shared 24 overlapping clones with the pre-treatment recurrent sample, 22 overlapping clones with the primary tumour, and 10 overlapping clones with the lymph node tumour, leaving 68% of its T-cell repertoire unique to the post-treatment recurrence. A similar effect of ICB was observed in HN386, from whom we collected only pre- and post-ICB recurrence samples: following ICB treatment, 45% of the post-ICB T-cell repertoire was not detected (unique) in the pre-ICB recurrent tumour despite sharing 106 overlapping clones (**Supplementary Table 3**, **Supplementary Figures 2G-H**). Together, these longitudinal cases suggest that tumour progression and therapeutic intervention reshape TCR dynamics through selection of pre-existing clones and expansion of newly recruited clones after ICB therapy.

These differences in clone diversity and sharing through disease stage motivated a closer examination of clonal expansion and their associations with cell states. We first categorized cells by clonotype size as unexpanded (1 cell), lowly expanded (2-10 cells), highly expanded (11-50 cells), or hyper-expanded (> 50 cells; **Figure 2G**). Within tumours, these expanded clonotypes typically reflect antigen-driven proliferation in response to tumour-associated antigens. Unsurprisingly, naïve and dysfunctional cells lie at the extremes of unexpanded and hyper-expansion, respectively (**Supplementary Figure I-J**). Central memory and effector memory cells showed intermediate levels of expansion. Proliferating cells were more frequently expanded, but less frequently hyperexpanded as dysfunctional cells. These patterns are consistent with a trajectory of antigen-driven activation, clonal expansion and eventual dysfunction due to chronic antigen stimulation in the tumour microenvironment.

To determine if these expanded populations were migrating across anatomically separated sites, we focused on patients with multi-sample TCR-profiling and labeled clones by whether they were detected in one or multiple samples. We termed these not-shared and shared clones, respectively (**Figure 2H**). In early disease, clones could be shared between primary tumour and TDLN. In intermediate and late disease, clones could be shared between any combination of Pri, LNMet, Rec, and Rec-pICB tumour tissues. Unexpanded clones were by definition never shared, and were also mostly enriched in the naïve T-cell population (**Figure 2I**). In contrast, shared expanded clones showed cell type distributions similar to all expanded clones, but were more prevalent in patients with late disease (**Figure 2J**). In these late disease patients, shared expanded clones made up an average of 12%% of each patient’s pan-sample clonotype repertoire (range 5.2-19%), with 2.4% (0.7-3.8%) belonging to highly or hyper-expanded clonotypes. These patterns of selective expansion and sharing of these specific T-cell states across distinct tumour sites suggest they are driven by shared antigen recognition, indicating that these shared-expanded clonotypes could be actively participating in tumour-associated reactivity.

## A signature of tumour-targeting T-cells

The selective expansion and cross-site sharing of these transcriptionally distinct clonotypes pointed to a common underlying driver of tumour antigen recognition. We therefore designated shared-expanded clonotypes as putative tumour-targeting CD8^⁺^ T cells and sought to define their molecular hallmarks. We extracted the transcriptomic and TCR profiles of 23,172 cells from patients with intermediate or late disease, partitioned the data into train and test sets using a 3:1 split (**Supplementary Figure 3A**), and performed initial exploration and model development using the training set only. Differential expression analysis (**Figure 3A**, **Supplementary Figure 3B-C**, **Supplementary Table 4**) between shared-expanded and not shared-expanded cells revealed 88 genes that were more highly abundant in shared-expanded cells (Log_2_FC>2, adjusted p>0.04). Conversely, 22 genes were more abundant in not shared-expanded cells (Log_2_FC<-2, adjusted p>0.04). These transcriptional differences between clonal expansion-motivated groups established the foundation for a machine learning classifier capable of identifying tumour-targeting CD8^⁺^ T cells without prior knowledge of tumour antigens. To build a robust model, we optimized multiple learners that each informed the final machine learning model (**Figure 3B, Methods**). Preliminary models included regularized generalized linear models (lasso, ridge, and elastic net) and a random forest. Input features for these preliminary models included global transcriptomics features and tissue origin, which codes whether the cell originated from a primary tumour, a lymph node metastasis, a distant recurrence, or post-pembrolizumab recurrence.

**Figure 3.**
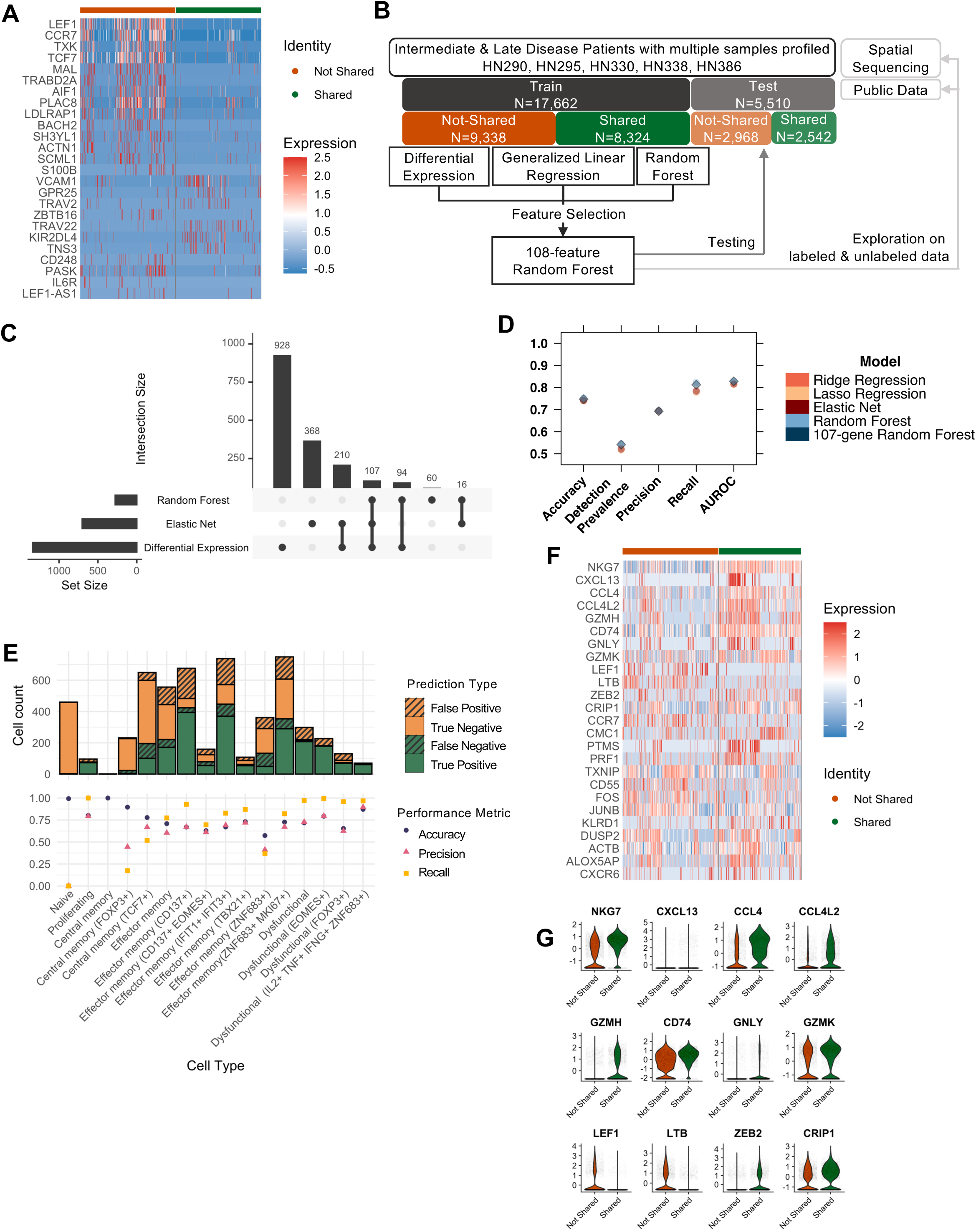
A gene expression signature to predict tumour reactive clones. A. Top 25 differentially expression genes between shared-expanded *vs.* not shared-expanded clones in the train data. Genes shown have FDR-adjusted p<0.05 and are ordered from largest fold change. B. Model training workflow for gene expression-based prediction of shared-expanded cells. Only cells from multi-sample intermediate or late disease patients were used in model training. The first round of model training fit generalized linear (elastic net, ridge, lasso regressions) and random forest preliminary models using the full feature set as input. Feature selection ranking from these models and differential expression results were used to select a final 108-feature set (107 genes + tissue-type). We trained a final 108-feature random forest using only the selected features and evaluated this signature on unlabeled, early disease, and public data. All models were evaluated using the hold-out test set. C. UpSet plot of feature overlap between elastic net regression (non-zero coefficients), random forest (top 25% cumulative variable importance), and differential expression analysis (adjusted p<0.05). Vertical bars show intersection sizes; horizontal bars show total features per method. The 107-gene consensus identified across all three methods, combined with tissue type as a covariate, defines the input feature set for the final 108-feature random forest classifier. D. Model performance metrics on the test set for all model types. Circles denote preliminary models; diamond denotes final random forest model. E. Final random forest model performance. Test set predictions by cell type, with top barplot showing breakdown of prediction type, and bottom dot plot showing performance metric summaries for accuracy (blue circle), precision (pink triangle), and recall (yellow square). F. Top 25 features by final 108-feature random forest variable importance in the test data. G. Distributions of top 12 features by final random forest variable importance in test data.

All preliminary models effectively predicted shared-expanded T-cells with a median accuracy of 80% (79-98%) and median train set AUC of 0.88 (0.87-0.99; **Supplementary Figure 3D**). To select a final feature set, we compared features between the best-performing generalized linear model (the elastic net), random forest model, and differentially expressed genes, (**Figure 3C**). These three complementary approaches converged on a common set of 107 genes that were consistently selected during model fitting and significantly differentially expressed. Incorporating tissue origin, which was also identified as an important feature by the machine learning models, resulted in a final set of 108 features.

With these 108-features, we optimized a final random forest model (108-feature RF) and tested all four models on the hold-out test set (**Figure 3D, Supplementary Figure 3D**). The 108-feature RF achieved comparable accuracy (73.7-74.7%, 108-feature RF: 74.7%), precision (67.8-69.4%, 108-feature RF: 69.2%) and recall (77.6-71.8%, 108-feature RF: 81.2%) compared with the full preliminary models, confirming that predictive information was concentrated within this reduced feature set. To evaluate cell type-specific prediction performance, we investigated precision and recall by each cell cluster (**Figure 3E, Supplementary Table 4**). The 108-feature RF correctly predicted and classified 99% (457/460) naïve cells as not shared-expanded and had similarly high precision (median 67%) and recall (median 83%) across cell groups. We observed modest performance reductions in specific populations of Central memory (FOXP3+), Central memory (TCF7+) and Effector memory (ZNF683+), likely due to their low relative abundance and transitory phenotype.

Finally, to determine whether the classifier had learned a transcriptional programme consistent with hallmarks of tumour reactivity, we examined the most important features as identified by the 108-feature RF. Comparing the top features between the test (**Figure 3F-G**) and train (**Supplementary Figure 3E-F**) sets showed consistent gene expression differences. Shared-expanded cells were enriched for genes associated with cytotoxicity (*NKG7*, *GZMK/H, PRF1, GNLY, CCL4*), and dysfunction *(CRIP1, ZEB2, KLRD1, CXCL13, PDCD1)*, indicating antigen-experienced T cells that have already undergone clonal expansion. In contrast, not shared-expanded cells were enriched for naïve and memory genes (*LEF1*, *LTB and CCR7*), characteristics of naïve T-cells that have yet to encounter antigen, as well as quiescent memory cells with limited expansion at the time of sampling (**Figure 3F-G, Supplementary Figure 3E-F**).

### Signature genes reflect known hallmarks of tumour reactivity and generalize to independent HNSCC datasets

Previous studies in lung^25^, melanoma^26^, gastrointestinal^27^ and pan-cancer solid tumours^28,29^ have identified distinct transcriptional profiles of tumour and viral reactivity in T-cells. To contextualize our signature within this broader landscape, we compiled five gene lists associated with tumour reactivity and three associated with viral reactivity from these published studies, and compared them with the 107 genes in our 108-feature RF (**Supplementary Figure 4A, Supplementary Table 5**). Of these 107 signature genes, 56 (52.3%) have been previously associated with tumour-reactivity in non-HNSCC tumours. These include top signature importance features such as *CXCL13* (described in all four studies), alongside *GNLY*, *LTB*, *TOX*, *ENTPD1*, *PDCD1*, *CD70*^30^ and *CXCR6*. These genes include established markers of chronic antigen exposure and T cell exhaustion, characteristics consistently attributed to tumour-reactive TILs, and have been implicated in tumour immune evasion. A further 29 signature genes (27.1%) were previously associated with viral reactivity in lung and melanoma, although five of these (*NKG7, CCL4, GZMH, GZMK, TCF7*) also have prior evidence of tumour reactivity, suggesting they reflect shared features of chronic antigen exposure rather than viral specificity alone. The biological relevance of these features, combined with the predictive performance of the 108-feature RF, suggested our classifier had learned a transcriptional programme consistent with known hallmarks of tumour reactivity.

To test whether this generalized beyond our study cohort, we next applied the classifier to an integrated atlas of 89,318 HNSCC CD8^⁺^ T cells. We obtained scRNA-seq data from seven published studies of HNSCC^22,31–36^, selecting CD8^⁺^ T cells by *CD8A*/*CD8B* expression and signature-based cell type annotation. We integrated these data together with our study data (**Supplementary Figure 4B**). Collectively, the data reflects CD8+ T cell profiles of HNSCC primary and metastatic tumours, pre-malignant lesions and normal tissues, and blood from both cancer and healthy patients (**Supplementary Figure 4C**). The signature accepts raw counts as input, assigns missing features as NA, and performs all necessary data transformations for model inference. Across the eight public datasets, there was a median overlap of 102 signature genes (range 101-107; **Supplementary Table 5**).

As no accompanying scTCR-seq data were available from these public datasets, definitive clonotype-based labels for tumour reactivity could not be derived. We therefore evaluated classifier performance *via* biological consistency, examining predicted tumour-reactive cells in the context of cell subtypes putatively associated with tumour specificity and across anatomical tissue distributions. Notably, our integrated UMAP projection comprising both our labeled study data and public data, showed close co-localization of predicted shared-expanded cells with our labeled shared-expanded cells (**Figure 4A**), indicating that the classifier identified transcriptionally concordant cells in independent data. Further investigation of top predictor genes showed consistent expression distributions in prediction-stratified groups between public datasets and our study data (**Supplementary Figure 4D**), providing evidence of cross-dataset transferability. Tumour tissue-specific cluster-level prediction breakdowns (**Figure 4B**) were also consistent across cell subtypes with the expected biology of tumour-reactive TILS: very few predicted shared-expanded cells were observed in naïve-like populations (median 0.33%, range 0-3.4%), and proportions were low in central memory (median 16%, 5.5-24%), and highest in proliferating (median 93%, 72—95%), cytotoxic (median 91%, 57-100%), and dysfunctional (median 90%, 83-97%) clusters (**Supplementary Table 5**).

**Figure 4.**
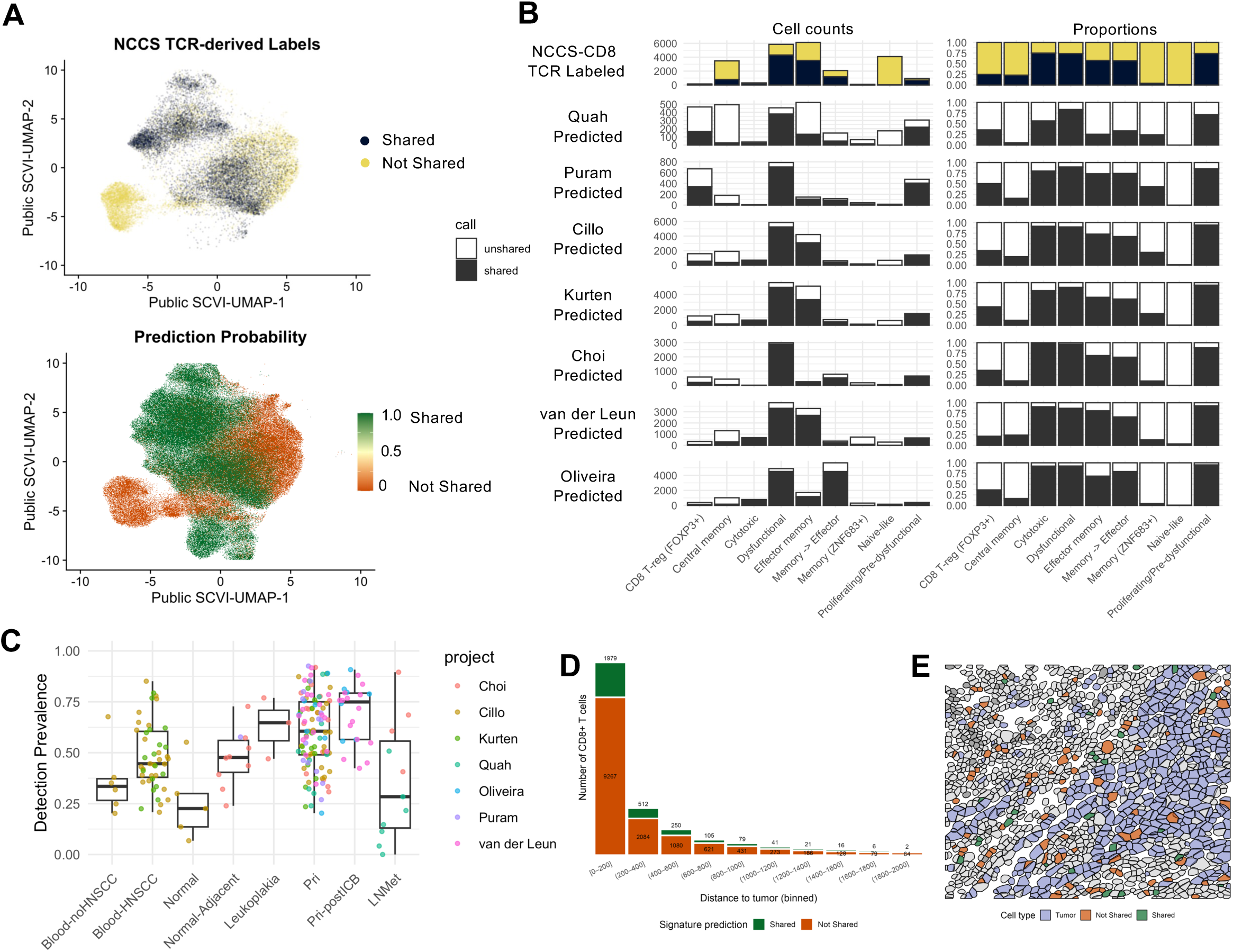
The tumour-reactive CD8⁺ T cell signature generalises across independent cohorts and resolves to tumour-proximal niches in situ. A. scTCR-seq-derived labels for shared-expanded (blue) *vs* not shared-expanded (yellow) in current study data subset from public HNSCC scRNA-seq integrated atlas (top), and 108-feature random forest signature predictions for public data based on downloaded scRNA-seq profiles (bottom). Prediction probabilities range from 100% (green) to 0% (orange). D. Breakdown of shared-expanded labels and classifier predictions across integrated atlas CD8^⁺^ T cell clusters. Upper panels show clonotype-based ground truth labels from the study cohort (shared-expanded, blue; not-shared-expanded, yellow). Lower panels show 108-feature RF predictions applied to all cells in the integrated atlas. Left panels show absolute cell counts; right panels show proportions. In each panel, full bar height represents the total cluster size; black bars indicate the shared-expanded or predicted shared-expanded count and proportion. B. Signature detection prevalence across tissue types in the integrated HNSCC CD8^⁺^ T cell atlas, calculated as the proportion of cells predicted shared-expanded by the 108-feature RF classifier. Each point represents one sample, coloured by dataset of origin. Box plots show the median (centre line), interquartile range (box limits), and 1.5× IQR (whiskers); points beyond the whiskers are outliers. C. Stacked bar plot of the number of CD8^⁺^ T cells across distance bins from the nearest tumor cell, stratified by shared and non-shared predicted signatures. D. Representative field of view reconstructed from the spatial transcriptomic data, showing the spatial distribution of tumor cells and CD8^⁺^ T cells classified as shared or non-shared.

The inclusion of multiple tissue types in the public data allowed us to explore signature predictions across peripheral blood, TDLN, and non-malignant, pre-malignant and malignant tissues (**Figure 4C, Supplementary Figure 4E-F**). Predicted shared-expanded proportions were lowest in non-cancer blood and normal tissues from healthy (included in Cillo *et al*^32^) and progressively higher in blood and tumour-adjacent normal tissues from HNSCC donors. Moreover, a consistent gradient of increasing prediction prevalence was observed across the clinical spectrum from tumour-adjacent normal to malignant tissue, independent of dataset-specific sources of variation. This increase in detection prevalence across increasingly malignant samples suggests our signature captures a biologically meaningful and progressive feature of the tumour immune microenvironment.

Collectively, the convergence of our signature genes with established transcriptional programmes of tumour reactivity, the consistent prediction patterns across seven independent public datasets, and the progressive enrichment of predicted tumour-reactive cells across the clinical spectrum support the robustness and generalisability of our classifier as a tool for identifying tumour-targeting CD8⁺ T cells in HNSCC.

### Spatial validation identifies tumour-proximal localization of predicted shared-expanded CD8+ T cells

A critical property of genuinely tumour-targeting T cells is their spatial relationship to malignant tissue: enrichment in tumour-proximal regions would provide orthogonal, spatially-resolved evidence of biological validity. We therefore applied the 108-feature random forest classifier to a spatial transcriptomics cohort comprising four HNSCC tissue sections profiled using a targeted 1,000-gene CosMx panel. Following preprocessing, batch correction, clustering, and cell type annotation, a total of 17,868 CD8+ T cells were identified across the four samples. Of the 107 genes included in the final classifier, 45 were also represented in the spatial panel. Missing features were assigned as NA prior to model inference, consistent with the approach used for public dataset validation.

Application of the classifier identified 3,015 CD8+ T cells (16.9%) as shared-expanded, and the remaining 14,853 cells as not shared-expanded. Predicted shared-expanded cells were detected across all tissue sections, although their abundance varied between samples. Proportions of predicted shared-expanded cells ranged from 7.8% in HN251_Pri and 8.1% in HN003_Pri (both primary tumours only) to 26.1% and 15.3% in HN279 primary tumour and matched, metastatic lymph node respectively. This demonstrates that our transcriptional signature associated with shared-expanded cells can also be applied across gene expression profiling platforms.

To investigate the spatial distribution of these cells, we calculated the Euclidean centroid-to-centroid distance between each CD8^+^ T cell and its nearest tumour cell and grouped cells into 200-pixel distance intervals. Across all samples, CD8^+^ T cells were most abundant in regions immediately adjacent to tumour cells and decreased progressively with increasing distance from tumour regions (**Figure 4D**). Predicted shared-expanded cells followed a similar pattern, with the largest number detected within 200 pixels of a tumour cell (n = 1,979), representing approximately 66% of all predicted shared-expanded cells. The abundance of predicted shared-expanded cells declined steadily with increasing distance from tumour cells.

Spatial reconstruction of representative fields of view demonstrated that predicted shared-expanded CD8+ T cells localized predominantly within tumour-rich regions whereas not shared-expanded cells were distributed more broadly throughout the surrounding tissue (**Figure 4E**). Together, these findings provide orthogonal spatial validation of the shared-expanded signature and demonstrate that the transcriptional programme identified from clonally shared and expanded T cells is associated with tumour-proximal localization within independent HNSCC specimens.

## Discussion

In this work, we set out to identify CD8+ T cells that recognize the tumour when the antigens they target are neither shared across patients nor tractable to define at scale. We reasoned that CD8^⁺^ T cells belonging to clones simultaneously expanded and shared across anatomical compartments and disease timepoints are enriched for cells responding to persistent tumour-associated epitopes and developed a transcriptomics signature to learn differences between cells belonging to TCR-labeled putatively tumour-reactive and non-tumour-reactive groups. Our 108-feature random forest signature was generalized across independent HNSCC cohorts, across tissue types spanning normal, pre-malignant and malignant states, and across profiling platforms, including a targeted FFPE spatial assay.

The genes that constitute this signature are not individually surprising. *CXCL13*, *TOX*, *ENTPD1*, *PDCD1*, *CXCR6*, together with cytotoxic effectors such as *NKG7*, *GZMK*, *GZMH* and *PRF1* recapitulate the chronic-antigen and dysfunction programme that has been defined through antigen-anchored approaches in melanoma, lung cancer, and pan-cancer epithelial tumours^25,26,29^. That two orthogonal strategies, antigen-resolved and clonal-behavioural, arrive at a shared transcriptional definition is evidence that our shared-expanded population is likely tumour-engaged, and extends the same reactive programme to HPV-negative oral cancer.

The main opposing hypothesis for these expanded, shared clones is that it is a bystander: a virus-specific or otherwise tumour-irrelevant T cell that expands and disseminates for reasons unconnected to the malignancy^5^. Two features of our study argue against bystanders dominating the signal. First, our cohort is uniformly HPV-negative, which removes the single largest source of intra-tumoural viral-antigen-specific CD8+ T cells in head and neck cancer. Some residual overlap we observe with viral-reactivity signatures is expected and most plausibly attributable to ubiquitous chronic antigen exposure rather than to an oncogenic virus resident within the tumour. Second, our antigen-agnostic signature independently recovered *ENTPD1* (CD39), a canonical marker discriminating tumour-reactive from bystander CD8^⁺^ T cells^5,28^. Convergence on CD39 through a purely clonal route is difficult to reconcile with a population dominated by bystanders.

Longitudinal sampling of patients across the evolution of their diseases allowed us to observe this population in motion. Shared-expanded clones were more prominent in primary tumours and recurrences than in draining lymph nodes, consistent with antigen-driven retention and progressive differentiation. This was also observed in recurrent tumour, months after ablation of the primary tumours and nodal stations. Most striking were the patients biopsied immediately before and shortly after initiating pembrolizumab, in whom the on-treatment repertoire was dominated by clones not detected before ICB. This pattern is consistent with clonal replacement as described following ICB in other epithelial and cutaneous cancers^20^. With only two treated patients, we advance this as a hypothesis to be tested in dedicated on-treatment cohorts in future work.

The most immediate value of this work is as a deployable biomarker of tumour-engaged CD8^⁺^ immunity in HPV-negative oral cancer, where archival formalin-fixed material is abundant but fresh tissue is less available. The signature retained discriminative behaviour on a targeted FFPE spatial platform using fewer than half its constituent genes, and predicted cells localized preferentially to tumour-proximal niches, suggesting that a tumour-targeting CD8^⁺^ readout can be recovered from routine pathological specimens and resolved *in situ*.

Several limitations qualify these conclusions. The signature was trained on a modest number of multi-sample patients with advanced disease, and broader validation incorporating antigen-resolved ground truth is required. The clonal expansion-based label necessarily weights the signature toward differentiated, antigen-experienced cells, with comparatively reduced sensitivity to the stem-like progenitor pool governing durability of anti-tumour responses^37,38^. More fundamentally, transcriptional inference remains a surrogate for the unsolved problem of predicting TCR-antigen-HLA recognition at scale. Until that ‘holy grail’ is computationally resolved, antigen-agnostic, clonally-grounded signatures such as this offer a practical route to identifying the tumour-reactive CD8^⁺^ T cells that shape immunotherapy responses in solid tumours.

The ability to identify tumour-targeting CD8^⁺^ T cells without knowledge of their cognate antigen addresses a fundamental bottleneck in cancer immunology. By leveraging clonal sharing and expansion as a proxy for tumour reactivity, and coupling this with single-cell transcriptomics across disease stages, tissue compartments, and treatment exposures, we define the molecular hallmarks of tumour-targeting CD8^⁺^ T cells in HPV-negative oral cancer and distil these into a transferable classifier. More broadly, the principle that clonal behaviour across space and time can serve as an antigen-agnostic proxy for tumour reactivity is likely generalizable beyond oral cancer, offering a transferable framework for mapping tumour-engaged immunity in solid tumours where the antigenic landscape remains undefined.

## Methods

### Ethics approval

This research complies with all relevant ethical regulations. The study of patient tumour samples was approved by SingHealth Centralized Institutional Review Board (CIRB: 2014/2093, 2018/2512 and 2016/2757) and each patient’s written consent.

### Patient sample collection

For tumours, all patients were confirmed histologically to be HNSCC and suitable for surgical resection (with no prior cancer treatment). Patients included males and females, aged 45–96. Only primary lesions larger than a T2 with sufficient tissue for study without compromising pathological exam were included. Details of clinical and pathologic features are provided in **Supplementary Table 1.** Fresh tissues were collected from the primary site as well as involved and non-involved tumour-draining lymph nodes, and transported to the laboratory within 30 min upon resection in the operating theater.

### Tumour sample dissociation

Tumours were minced and transferred into C Tubes, then digested using the Human Tumour Dissociation Kit and gentleMACS Octo Dissociator as described previously and in the manufacturer’s protocol (all from Miltenyi Biotech)^22^. After digestion, single-cell suspensions were passed through 70µM cell strainers (VWR), washed with RPMI supplemented with 10% FBS, and pelleted by centrifugation at 300 x g for 5 minutes. Red blood cells were lysed when necessary to remove erythrocyte contamination. Subsequently, cells were resuspended in appropriate medium and counted using the Countess 3 Automated Cell Counter (Invitrogen) with trypan blue for dead cell exclusion.

### Cell enrichment by magnetic separation

No more than 1×10^7^ of single cell suspensions from each tissue were magnetically labelled with anti-CD8 (Miltenyi Biotech), except for samples HN307_Pri, HN307_sLN, HN309_Pri and HN309_sLN which were labelled with anti-CD45 microbeads (Miltenyi Biotech). Labelling was performed in MACS buffer (0.5% BSA and 2mM EDTA in PBS), and incubated for 20mins at 4°C. After incubation, cells were washed with MACS buffer, filtered through 40µM cell strainers (VWR), pelleted and resuspended in 500µL of MACS buffer prior to magnetic separation using MS column and a magnetic stand, as described in the manufacturer’s protocol. Following separation, cells were washed, pelleted and resuspended in 0.04% BSA in PBS for cell counting and single-cell capture.

### Single-cell RNA- and TCR-sequencing

Single-cell 5’ gene expression (GEX) and TCR libraries were prepared using the Chromium Next GEM Single Cell 5’ Reagent Kits v2 (10x Genomics), according to the manufacturer’s protocol. Freshly sorted cells were resuspended in 0.04% BSA in PBS and adjusted to a final concentration of 500-1200 cells/µL. Subsequently, cells and reagents were loaded into Chromium Next GEM Chip K for gel bead-in emulsion (GEM) generation and barcoding, targeting a recovery of 2000-8000 cells per sample. Within each GEM, RNA was reverse transcribed, followed by GEM breakage and cDNA purification using Dynabeads MyOne SILANE (included in the kit). The resulting cDNA was amplified and used for the construction of both GEX and TCR libraries. For GEX library construction, 2–50 ng of amplified cDNA was fragmented, end repaired and size-selected using SPRIselect Reagent (Beckman Coulter), followed by sample index PCR. For TCR library construction, 2 µl of amplified cDNA was enriched for V(D)J sequence using the Human T Cell V(D)J Enrichment Kit (10x Genomics). Subsequently, 2–50 ng of the enriched transcripts was fragmented, end repaired, sample indexed and size-selected using SPRIselect Reagent (Beckman Coulter). Final libraries were sequenced using an Illumina NovaSeq 6000 seq (Illumina) with 150 bp paired-end reads.

### scRNAseq data pre-processing and integration

After sequencing, single-cell RNA-seq (scRNA-seq) reads were aligned to the GRCh38-2020-A reference transcriptome and quantified using Cellranger count (v7.1.0, 10x Genomics). Raw gene expression data processed using Seurat (v5.0.0)^39^ and cells were filtered for quality using sample specific cutoffs (minimal gene counts > 500, minimal UMI count > 200, and maximal mitochondrial fraction < 8%). Highly variable genes (HVGs) were identified using sc.pp.highly_variable_genes() with the cell_ranger flavour after normalization and log-transformation.

Integration was performed using the conditional variational autoencoder implemented in scVI^40^. The scVI model was set up on raw counts with patient identity as the batch key. The number of training epochs was determined adaptively using the heuristic max_epochs = min(round((20000 / n_cells) × 400), 400). Integration quality was assessed by visual inspection of UMAP projections coloured by patient identity and tissue-level covariables including sample type, specimen type, tissue type, mitochondrial fraction, ribosomal fraction, and total counts, confirming no residual structure attributable to technical or patient-level variation after correction.

### CD8+ T cell filtering, clustering, and annotation

CD8+ T cells were identified by filtering on CD8A/CD8B expression and signature-based cell type annotation using CellTypist^41^ and Azimuth^42^ to obtain a high-confidence population of 51,459 CD8^+^ T-Cells across the 28 samples (**Figure 1B**). CD8^+^ subset cell clusters were assigned using a two-stage Louvain clustering approach. First, a coarse resolution of 0.2 was used to cluster cells into 5 major clusters, each representing canonical CD8^+^ T cell states (**Supplementary Fig 1**). Major clusters were subclustered independently to resolve fine-grained heterogeneity, identifying seven effector memory, three central memory, and three dysfunctional subpopulations. Final cell type labels and defining marker genes are provided in **Supplementary Table 2**. Cell cluster identities were assigned based on the following marker combinations: Naïve (*CCR7+, LEF1+, IL7R+*), Proliferating (*MKI67+*), Dysfunctional (*CXCL13+, LAG3+, PDCD1+*), Central memory (*IL7R+, CCR7+, GZMK+*), Effector memory (*IL7R+, CCR7-, GZMK+*).

### scTCRseq data pre-processing

Single-cell TCR-seq (scTCR-seq) reads were aligned to the vdj_GRCh38_alts_ensembl-7.1.0 V(D)J reference and quantified using cellranger vdj (v7.1.0 10x Genomics), producing filtered contig annotation files for each sample. TCR data were loaded and integrated with gene expression data into a joint MuData object using Scirpy^43^ (v0.17.2) and Muon (v0.1.6). Cell-level overlap between modalities was assessed per sample by comparing barcode intersections. Chain quality control was performed using Scirpy’s built-in chain indexing and QC functions, and cells with more than two productive TCR chain pairs (multichain cells) were removed prior to clonotype assignment.

### TCR repertoire analysis

TCR contig data were loaded from Cell Ranger V(D)J filtered contig annotation files for each sample and processed using two complementary pipelines. Initial clonotype definition and shared-expanded labelling were performed in Python using Scirpy (version 0.17.2), integrated within the scVI-based gene expression analysis pipeline. To enable joint analysis with the Seurat-based gene expression object and to leverage scRepertoire’s repertoire analysis functionality, TCR data were subsequently reprocessed in R using scRepertoire^44^ (v 2.5.8). Cell-level overlap between TCR and gene expression modalities was confirmed per sample by barcode intersection.

A clonotype was defined by the exact paired CDR3 nucleotide sequence of both the α and β chains (CTnt), consistent with the nucleotide identity-based clonotype definition applied in the Scirpy pipeline. Clonotype size was calculated per clone and used to categorize cells as unexpanded (1 cell), lowly expanded (2–10 cells), highly expanded (11–50 cells), or hyperexpanded (>50 cells). Cross-sample clonal overlap was assessed using clonalCompare().

A clonotype was defined as shared if the identical paired CDR3 nucleotide sequence was detected in more than one sample from the same individual. A clonotype was defined as shared-expanded if it was both shared across samples and clonally expanded (≥2 cells). All shared-expanded clones are therefore always expanded, by definition. These binary labels were transferred to the Seurat object metadata and used as the primary class labels for all downstream differential expression and classifier development analyses. Consistency between shared-expanded labels derived from scRepertoire and those derived from the Scirpy pipeline was confirmed by cross-tabulation of barcode-matched labels across both tools.

### Data partitioning

We started with 23,172 cells with both scTCR-seq and scRNA-seq data from multi-sample patients with intermediate (*ie*. local metastatic: HN290) and late (*ie*. distant metastatic disease: HN295, HN330, HN338, HN386). We split them into 75% train and 25% test sets with balancing for shared-expansion (our label of interest), tissue type and cell type. These partitions were maintained in differential expression analysis and throughout model training.

### Differential expression analysis

Differential expression between shared-expanded and not-shared-expanded CD8^⁺^ T cells in the train partition was performed using logistic regression as implemented in Seurat’s FindMarkers() function (v 5.0.0). Patient identity and tissue type were included as latent variables to account for inter-patient heterogeneity and tissue-specific transcriptional variation, both of which could confound the comparison between clonally defined groups. Genes were considered differentially expressed at an absolute log2 fold-change threshold of >2 and an adjusted p-value of <0.05 after Benjamini-Hochberg correction. Differentially expressed genes were used as one of three complementary inputs to feature selection for classifier development. Results of train-set differential expression analysis were investigated in the test set.

### Machine learning signature development

All model training and evaluation was performed in R using the tidymodels framework (version 1.4.1). Gene expression input features were extracted as raw counts, and tissue type was encoded as an ordered factor with levels: primary tumour (Pri), lymph node (LN), recurrence (Rec), and post-ICB recurrence (Rec-pICB). A preprocessing recipe was defined using the recipes package (v 1.3.1), comprising the following sequential steps: median imputation of missing numeric predictors (step_impute_median()); removal of zero-variance predictors (step_zv()); normalisation of all numeric predictors (step_normalize()); handling of novel tissue type levels unseen during training (step_novel()); encoding of missing tissue type values as unknown (step_unknown()); and one-hot encoding of tissue type as dummy variables (step_dummy()).

Four preliminary models were trained on the full transcriptional feature set plus tissue type using 10-fold cross-validation on the training set. Generalized linear models were implementing using the glmnet engine (v 4.1.10). For elastic net-regularized logistic regression, both the penalty (log10 scale, range −2 to 2) and mixture parameters were tuned over a regular grid of 50 levels each. Lasso and ridge models were tuned over the penalty parameter alone. A random forest classifier was implemented using the ranger engine (v0.18.0) with 1,000 trees. The number of randomly selected predictors at each split (mtry, range 10–150) and the minimum number of data points required for a node to be split (min_n, range 10–40) were tuned over a regular grid of 5 levels each. All hyperparameter tuning was performed in parallel using the doParallel package (v1.0.17). The best-performing hyperparameter combination for each model was selected by AUC on the cross-validation resamples.

A final feature set of 107 genes was derived from the intersection of features consistently selected by the elastic net, the random forest variable importance ranking, and the differentially expressed gene list. Tissue type was additionally included, yielding a final 108-feature set. The identical preprocessing recipe was applied to the reduced feature set and a final random forest model (108-feature RF) was trained using the same hyperparameter tuning procedure described above. The final model was evaluated on the held-out test set and performance was assessed by accuracy, precision, recall, and AUC. Variable importance was quantified using mean decrease in Gini impurity.

### Public dataset integration and external validation

Processed count matrices from seven published HNSCC scRNA-seq studies were obtained from GEO accessions <u>GSE164690</u>^36^, <u>GSE181919</u>^31^, <u>GSE232240</u>^35^, <u>GSE182227</u>^34^, <u>GSE139324</u>^32^ and <u>GSE188737</u>^22^. Raw scRNA-seq data from Oliveira *et al.*^33^ can be accessed at dbGaP accession <u>phs002864.v1.p1</u>. The data was checked for duplicate samples. CD8+ T cells were isolated by filtering on CD8A and CD8B expression and signature-based cell type annotation, then integrated with our study data using the same preprocessing and batch correction pipeline described above. The 108-feature RF was applied to each public dataset independently using raw counts as input; genes absent from a given dataset were assigned as NA prior to inference. Tissue type values were re-coded to the closest match to Pri/LNMet/TDLN/Rec, but otherwise omitted (*eg.* normal blood treated as missing value). In the absence of clonotype-based ground truth labels, classifier performance was evaluated through biological consistency across annotated cell subtypes and tissue types.

### Spatial transcriptomics

Formalin-fixed paraffin-embedded (FFPE) tumour tissue sections were prepared for profiling using the CosMx Spatial Molecular Imager (SMI; Bruker) according to the manufacturer’s protocol. Briefly, 5 µm-thick tissue sections were mounted on VWR Superfrost Plus Micro Slides (VWR), baked at 37°C overnight and stored at 4°C. Tissue sections underwent standard deparaffinization, target retrieval and enzymatic digestion prior to loading into the instrument for downstream profiling. Spatial transcriptomics was performed using the CosMx Human Universal Cell Characterization RNA Panel (1000-plex) together with morphology markers (B2M/CD298, PanCK, CD45, CD68 antibodies) and DNA staining (DAPI). Image processing, cell segmentation, transcript assignment to cells and generation of count matrix were performed using the AtoMx Spatial Informatics Platform (SIP). Cell boundaries were mapped using the multi-modal CosMx segmentation pipeline by levering both nuclear and membrane stains.

### Statistical Analysis & Data Visualization

All statistical analyses and data visualisation were performed in R (v4.5.1) and Python (v3.9.16) using the packages described above. Data visualizations were generated in R using BPG^45^ (v5.9.8), ggplot2 (v4.0.0) and patchwork (v 1.3.2).

Non-parametric tests were used for group comparisons unless otherwise stated. Multiple testing correction was applied using the Benjamini-Hochberg method. Statistical significance was defined using a p-value or adjusted p-value threshold of 0.05.

## Data and code availability

Raw and processed single-cell RNA sequencing and T cell receptor sequencing data generated in this study have been deposited in the Gene Expression Omnibus (GEO) under accession GSE345548 and will be made publicly available upon publication. Spatial transcriptomics data generated in this study have been deposited under GSE345753 and will be made publicly available upon publication. The integrated HNSCC CD8^⁺^ T cell atlas, comprising data from this study and six published public datasets, omitting access-protected data from Oliveira *et al*^33^, is available at DOI: 10.5281/zenodo.22258683. Original public datasets reanalysed in this study are available under the accessions <u>GSE164690</u>^36^, <u>GSE181919</u>^31^, <u>GSE232240</u>^35^, <u>GSE182227</u>^34^, <u>GSE139324</u>^32^ <u>GSE188737</u>^22^ and <u>phs002864.v1.p1</u>^33^.

Code to produce results reported in this manuscript, the 108-feature random forest classifier, and associated code to run the signature on new data are available at https://github.com/nccsCancerTherapeuticsLab/HNSCC_CD8_TumourReactivity, and archived at DOI: 10.5281/zenodo.22258683.

## Acknowledgments

We thank all patients and families who generously contributed to this project. Access for 10x experiments were provided by the Laboratory of Cell Therapy and Cancer Vaccine, National Cancer Center Singapore (under Dr Han-Chong Toh). The computational work for this article was partially performed on resources of the National Supercomputing Centre, Singapore (https://www.nscc.sg). This project was funded through the following grants to NGI, for which we are truly grateful: National Medical Research Council (Singapore) STAR (MOH-001684-00) and Individual Research Grants (MOH-001701-00), and the Peter Fu Head and Neck Cancer Program (under the Oncology Academic Clinical Program, National Cancer Centre Singapore).

## Author Contributions

CHL, HSQ, CA, and NGI conceptualized and designed the study, and interpreted the data. HSQ, CA, LS, HB, HSL, FTC and DT performed the experiments, and were supervised by SW, SKB and NGI. CHL carried out bioinformatic analyses supported by CA. CHL and HSQ drafted the manuscript, critically revised and approved the final version together with NGI.

## References

1. Tumeh, P. C. et al. PD-1 blockade induces responses by inhibiting adaptive immune resistance. Nature 515, 568–571 (2014).

2. Klein, C., Brinkmann, U., Reichert, J. M. & Kontermann, R. E. The present and future of bispecific antibodies for cancer therapy. Nat. Rev. Drug Discov. 23, 301–319 (2024).

3. Rafei, H., Upadhyay, R. & Sharma, P. A guide to CAR T cell therapies: development, current status and future prospects. Nat. Rev. Immunol. 10.1038/s41577-026-01322-1 (2026) doi:10.1038/s41577-026-01322-1.

4. Rosenberg, S. A. & Restifo, N. P. Adoptive cell transfer as personalized immunotherapy for human cancer. Science 348, 62–68 (2015).

5. Simoni, Y. et al. Bystander CD8+ T cells are abundant and phenotypically distinct in human tumour infiltrates. Nature 557, 575–579 (2018).

6. Scheper, W. et al. Low and variable tumor reactivity of the intratumoral TCR repertoire in human cancers. Nat. Med. 25, 89–94 (2019).

7. Arnaud, M. et al. Sensitive identification of neoantigens and cognate TCRs in human solid tumors. Nat. Biotechnol. 40, 656–660 (2022).

8. Tippalagama, R. et al. Antigen-specificity measurements are the key to understanding T cell responses. Front. Immunol. 14, 1127470 (2023).

9. Draper, L. M. et al. Targeting of HPV-16+ Epithelial Cancer Cells by TCR Gene Engineered T Cells Directed against E6. Clin. Cancer Res. Off. J. Am. Assoc. Cancer Res. 21, 4431–4439 (2015).

10. Johnson, L. A. et al. Gene therapy with human and mouse T-cell receptors mediates cancer regression and targets normal tissues expressing cognate antigen. Blood 114, 535–546 (2009).

11. Leidner, R. et al. Neoantigen T-Cell Receptor Gene Therapy in Pancreatic Cancer. N. Engl. J. Med. 386, 2112–2119 (2022).

12. Dunn, G. P., Bruce, A. T., Ikeda, H., Old, L. J. & Schreiber, R. D. Cancer immunoediting: from immunosurveillance to tumor escape. Nat. Immunol. 3, 991–998 (2002).

13. Schreiber, R. D., Old, L. J. & Smyth, M. J. Cancer immunoediting: integrating immunity’s roles in cancer suppression and promotion. Science 331, 1565–1570 (2011).

14. Liu, X., Harbison, R. A., Varvares, M. A., Puram, S. V. & Peng, G. Immunotherapeutic strategies in head and neck cancer: challenges and opportunities. J. Clin. Invest. 135, e188128 (2025).

15. Schumacher, T. N. & Schreiber, R. D. Neoantigens in cancer immunotherapy. Science 348, 69–74 (2015).

16. Lybaert, L. et al. Challenges in neoantigen-directed therapeutics. Cancer Cell 41, 15–40 (2023).

17. O’Brien, H. et al. Breaking the performance ceiling for neoantigen immunogenicity prediction. Nat. Cancer 4, 1618–1621 (2023).

18. Oliveira, G. & Wu, C. J. Dynamics and specificities of T cells in cancer immunotherapy. Nat. Rev. Cancer 23, 295–316 (2023).

19. Zheng, L. et al. Pan-cancer single-cell landscape of tumor-infiltrating T cells. Science 374, abe6474 (2021).

20. Yost, K. E. et al. Clonal replacement of tumor-specific T cells following PD-1 blockade. Nat. Med. 25, 1251–1259 (2019).

21. Shorer, O., Pinhasi, A. & Yizhak, K. Single-cell meta-analysis of T cells reveals clonal dynamics of response to checkpoint immunotherapy. Cell Genomics 5, 100842 (2025).

22. Quah, H. S. et al. Single cell analysis in head and neck cancer reveals potential immune evasion mechanisms during early metastasis. Nat. Commun. 14, 1680 (2023).

23. Wolfl, M. et al. Activation-induced expression of CD137 permits detection, isolation, and expansion of the full repertoire of CD8+ T cells responding to antigen without requiring knowledge of epitope specificities. Blood 110, 201–210 (2007).

24. Connolly, K. A. et al. A reservoir of stem-like CD8+ T cells in the tumor-draining lymph node preserves the ongoing antitumor immune response. Sci. Immunol. 6, eabg7836 (2021).

25. Caushi, J. X. et al. Transcriptional programs of neoantigen-specific TIL in anti-PD-1-treated lung cancers. Nature 596, 126–132 (2021).

26. Oliveira, G. et al. Phenotype, specificity and avidity of antitumour CD8+ T cells in melanoma. Nature 596, 119–125 (2021).

27. Zheng, C. et al. Transcriptomic profiles of neoantigen-reactive T cells in human gastrointestinal cancers. Cancer Cell 40, 410–423.e7 (2022).

28. Duhen, T. et al. Co-expression of CD39 and CD103 identifies tumor-reactive CD8 T cells in human solid tumors. Nat. Commun. 9, 2724 (2018).

29. Lowery, F. J. et al. Molecular signatures of antitumor neoantigen-reactive T cells from metastatic human cancers. Science 375, 877–884 (2022).

30. Huang, J. et al. Modulation by IL-2 of CD70 and CD27 expression on CD8+ T cells: importance for the therapeutic effectiveness of cell transfer immunotherapy. J. Immunol. 176, 7726–7735 (2006).

31. Choi, J.-H. et al. Single-cell transcriptome profiling of the stepwise progression of head and neck cancer. Nat. Commun. 14, 1055 (2023).

32. Cillo, A. R. et al. Immune Landscape of Viral- and Carcinogen-Driven Head and Neck Cancer. Immunity 52, 183–199.e9 (2020).

33. Oliveira, G. et al. Preexisting tumor-resident T cells with cytotoxic potential associate with response to neoadjuvant anti-PD-1 in head and neck cancer. Sci. Immunol. 8, eadf4968 (2023).

34. Puram, S. V. et al. Cellular states are coupled to genomic and viral heterogeneity in HPV-related oropharyngeal carcinoma. Nat. Genet. 55, 640–650 (2023).

35. Van Der Leun, A. M. et al. Dual Immune Checkpoint Blockade Induces Analogous Alterations in the Dysfunctional CD8+ T-cell and Activated Treg Compartment. Cancer Discov. 13, 2212–2227 (2023).

36. Kürten, C. H. L. et al. Investigating immune and non-immune cell interactions in head and neck tumors by single-cell RNA sequencing. Nat. Commun. 12, 7338 (2021).

37. Krishna, S. et al. Stem-like CD8 T cells mediate response of adoptive cell immunotherapy against human cancer. Science 370, 1328–1334 (2020).

38. Siddiqui, I. et al. Intratumoral Tcf1+PD-1+CD8+ T Cells with Stem-like Properties Promote Tumor Control in Response to Vaccination and Checkpoint Blockade Immunotherapy. Immunity 50, 195–211.e10 (2019).

39. Hao, Y. et al. Dictionary learning for integrative, multimodal and scalable single-cell analysis. Nat. Biotechnol. 42, 293–304 (2024).

40. Lopez, R., Regier, J., Cole, M. B., Jordan, M. I. & Yosef, N. Deep generative modeling for single-cell transcriptomics. Nat. Methods 15, 1053–1058 (2018).

41. Domínguez Conde, C. et al. Cross-tissue immune cell analysis reveals tissue-specific features in humans. Science 376, eabl5197 (2022).

42. Butler, A. et al. Azimuth: A Shiny App Demonstrating a Query-Reference Mapping Algorithm for Single-Cell Data. (2023).

43. Sturm, G. et al. Scirpy: a Scanpy extension for analyzing single-cell T-cell receptor-sequencing data. Bioinformatics 36, 4817–4818 (2020).

44. Borcherding, N., Bormann, N. L. & Kraus, G. scRepertoire: An R-based toolkit for single-cell immune receptor analysis. F1000Research 9, 47 (2020).

45. P’ng, C. et al. BPG: Seamless, automated and interactive visualization of scientific data. BMC Bioinformatics 20, (2019).

