## Supplementary figures and images for "Integrative single cell analysis of CD8+ T-cells across early and advanced oral cancers reveals signatures of anti-tumour activity"

### Supplemental Figures 1-4

#### Supplementary Figure 1

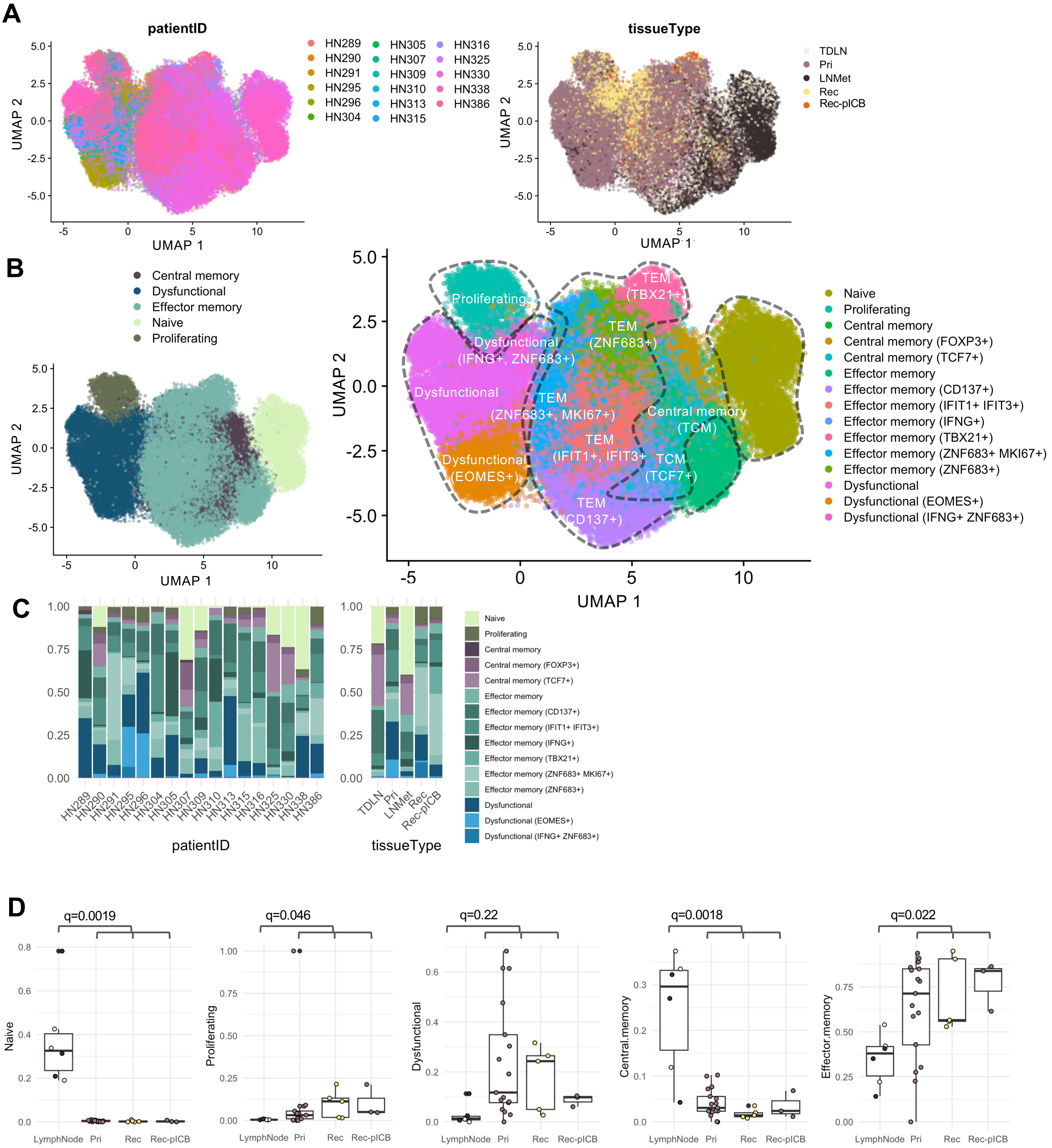

#### Supplementary Figure 2

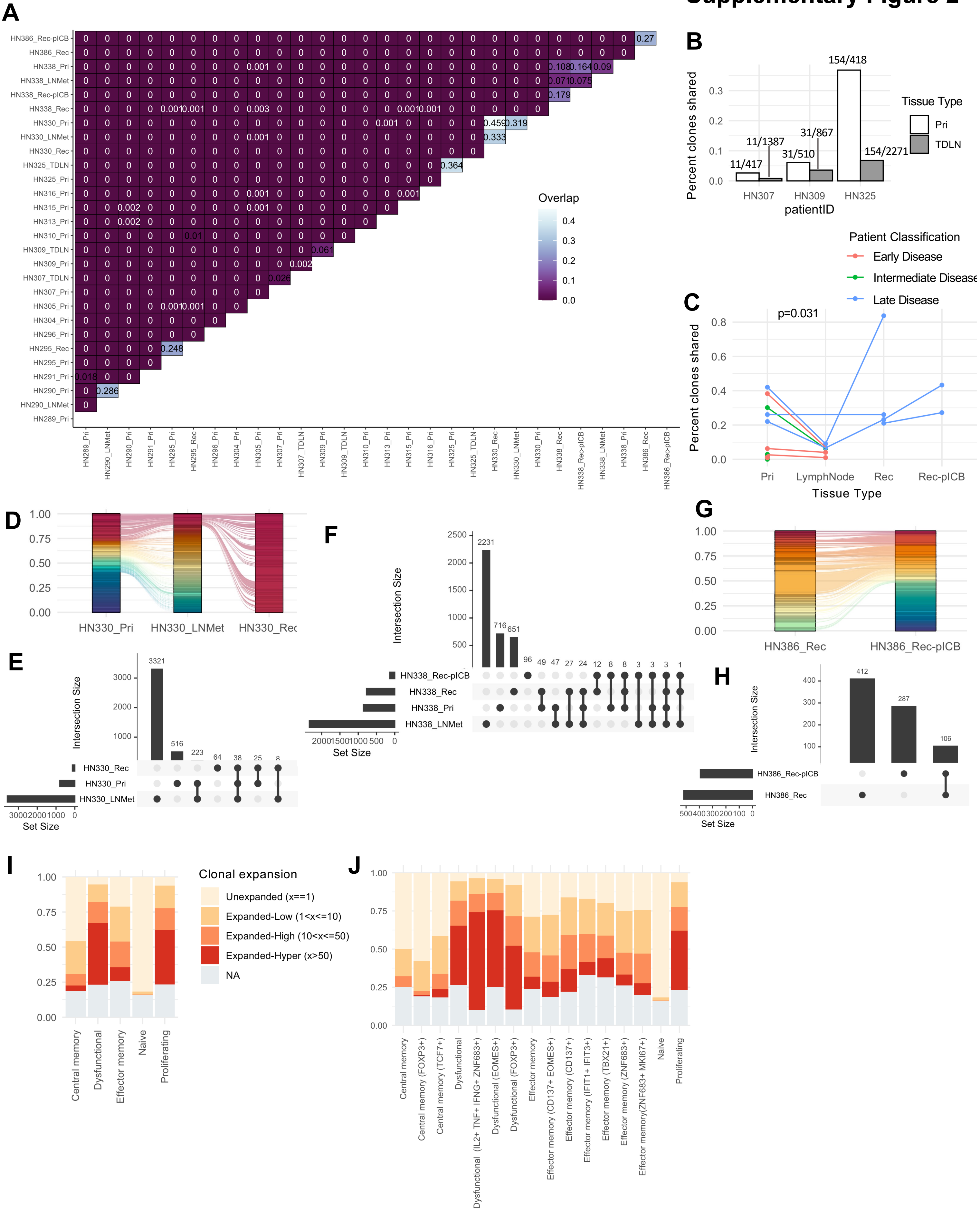

#### Supplementary Figure 3

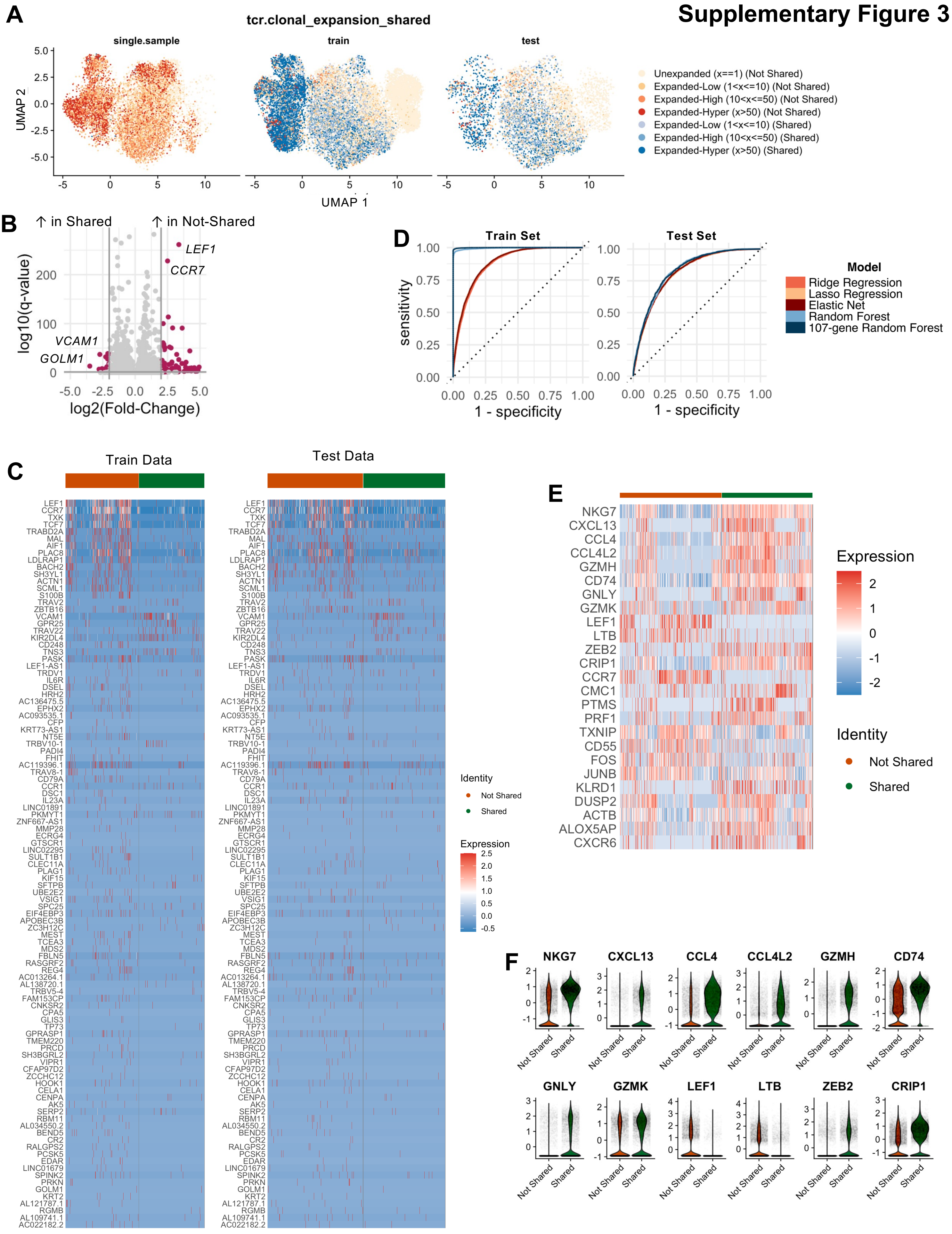

#### Supplementary Figure 4

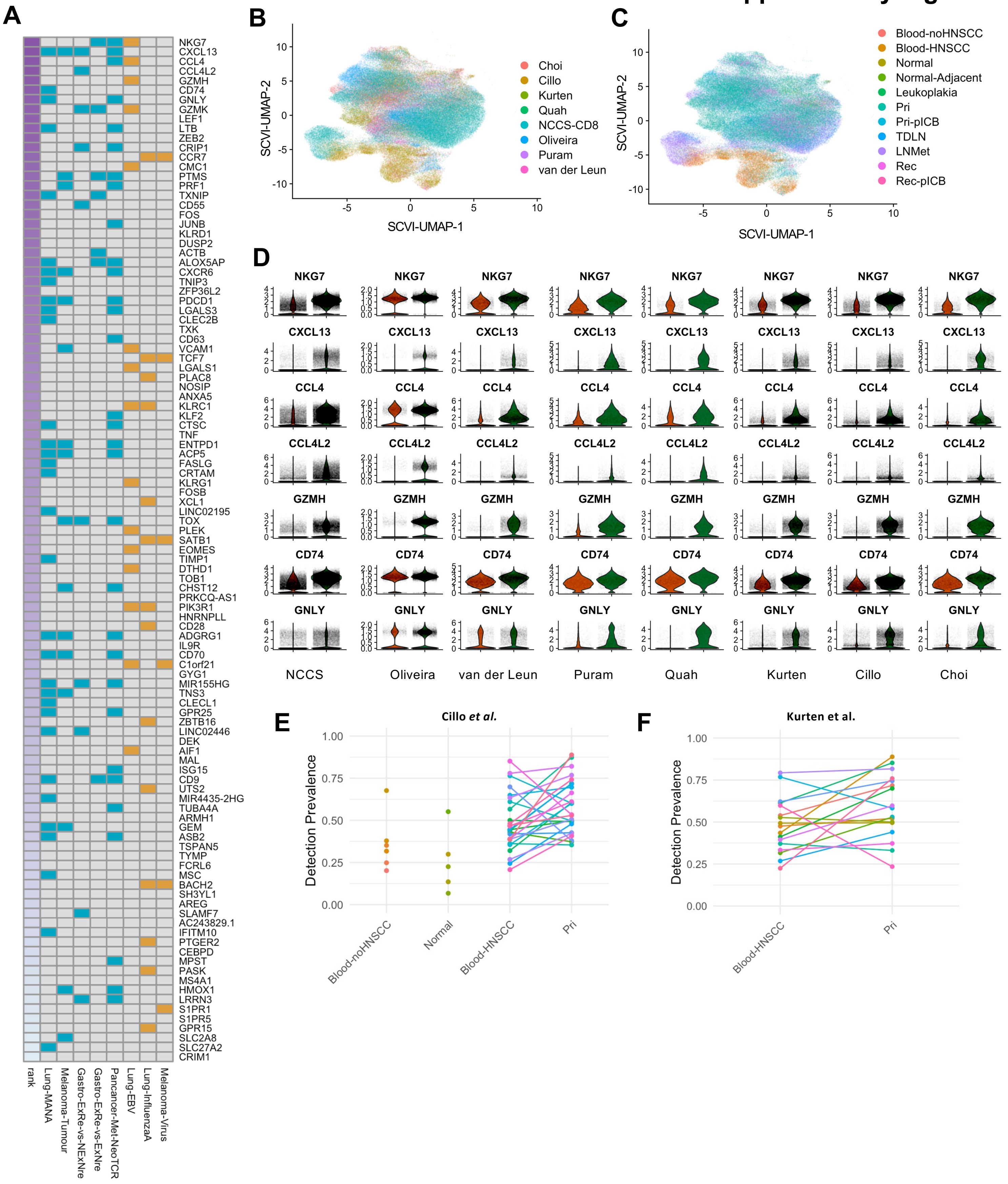
